# Aquila_stLFR: diploid genome assembly based structural variant calling package for stLFR linked-read

**DOI:** 10.1101/742239

**Authors:** Yichen Henry Liu, Griffin L. Grubbs, Lu Zhang, Xiaodong Fang, David L. Dill, Arend Sidow, Xin Zhou

## Abstract

**Motivation:** Identifying structural variants (SVs) is of critical importance in health and disease, however, detecting them remains a scientific and computing challenge. Several linked-read sequencing technologies, including 10X linked-read, TELL-Seq, and single tube long fragment read (stLFR), have been recently developed as cost-effective approaches to reconstruct multi-megabase haplotypes (phase blocks) from sequence data of a single sample. These technologies provide an optimal sequencing platform to characterize SVs, though few computational algorithms can utilize them. Thus, we developed Aquila_stLFR, an approach that resolves SVs through haplotype-based assembly of stLFR linked-reads.

**Results:** Aquila_stLFR first partitions LFRs into two haplotype-specific blocks, by taking advantage of the potential phasing ability of the linked-read itself. Each haplotype is then assembled independently, to achieve a complete diploid assembly to finally reconstruct the genome-wide SVs. We benchmarked Aquila_stLFR on a well-studied sample, NA24385, and showed Aquila_stLFR can detect medium to large size (50bp – 10kb) deletions with a high sensitivity and insertions with a high specificity.

**Availability:** Source code and documentation are available on https://github.com/maiziex/Aquila_stLFR.

**Supplementary information:** Supplementary data are available at *Bioinformatics* online.

## 1 Introduction

Generating a precise and customized diploid human genome for each individual will be a breakthrough for uncovering the fundamental relationship between genotype and phenotype, having far reaching health implications, e.g. in cancer-related variants and risk for genetic disease (Tewhey *et al*., 2011). Illumina short-read sequencing has had a major influence on human genetic studies. Over 100,000 individual genomes have been sequenced, allowing detection of unique, disease-causing variations in personal genomes (Lunshof *et al*., 2010). Large-scale genome studies, such as the 1000 Genome Project and the 10k UK Genome Project, have relied on reference-based alignment approaches, making great progress in uncovering genomic differences among individuals. However, identifying individual variations in highly variable or repetitive regions has been less accurate due to limitations of the resequencing technology (Sohn *et al*., 2018). Structural variants (SVs ≥ 50bp), which have shown a significant phenotypic impact by disrupting gene function and regulation, are also challenging to detect through short reads by reference-based alignment algorithms. *De novo* assembly is a better alternative for building a precise diploid genome on a large scale to identify SVs from haploid assemblies. To date, different assembly-based methods have been developed to offer a powerful approach to identify SVs (Wala *et al*., 2018; Fan *et al*., 2017; Nattestad *et al*., 2016), even though the breakpoints of large variants are less likely to be spanned through short reads.

The recently developed 10X linked-read, TELL-Seq, and single tube long fragment read (stLFR) sequencing technologies offer cost-effective solutions for large-scale “perfect genome” assembly (Peters *et al*., 2014; Zheng *et al*., 2016; McElwain *et al*., 2017) that enable optimal sequencing platforms to better characterize SVs than short reads. Compared to short-read and long-read technologies, these linked-read technologies combine low sequencing error and long-range contiguity. stLFR enables co-barcoding of over 8 million 20 – 300 kb genomic DNA fragments, and these long-range fragments enable efficient phasing of variants, resulting in long phase block N50 (34MB for NA12878, Wang *et al*., 2019). The long-range information of 10X linked-read allows detection of SVs, *de novo* mutations, and haplotype phasing much easier and accurately (Zheng *et al*., 2016; Zhou *et al*., 2018). Taking full advantage of these linked-read data to generate a diploid assembly and detect SVs requires development of new algorithms. Supernova was introduced by 10X Genomics to assemble 10X linked-read sequencing data (Weisenfeld *et al*., 2017), though its performance has limitations in identifying SVs from diploid assembly (Zhang *et al*., 2019, 2020; Zhou *et al*., 2021). Another software, Aquila, was developed recently to solve this problem for 10X linked-reads by achieving diploid assemblies and reconstructing genome-wide SVs with high sensitivity and accuracy (Zhou *et al*., 2021). Here, we develop Aquila_stLFR, which extends Aquila to adapt to the key characteristics of stLFR to identify SVs from diploid assembly with a higher accuracy compared with other state-of-the-art methods. Furthermore, we introduce a hybrid assembly mode, “Aquila_hybrid”, to enable combining of both stLFR and 10X linked-read data.

## 2 Methods

Aquila_stLFR is a reference-assisted, local *de novo* assembly pipeline (Supplementary Figure 1). The reference is used to allocate long fragment reads (LFRs) into haplotype-specific blocks, and local assembly is then performed within small chunks independently for both haplotype-specific blocks. The input files for Aquila_stLFR are aFASTQ file with paired short reads, a BAM file, and a VCF file (by FreeBayes, Garrison *et al*., 2012). To generate the BAM file through BWA-MEM (Li *et al*., 2009), each sequence identifier line in the FASTQ file needs to contain the barcode identifier, starting with “BX:Z:” (for instance, “BX:Z:540_839_548” where 540_839_548 is the barcode identifier, Supplementary Figure 2). The BAM file by BWA-MEM will then also include the extra field “BX:Z:” for Aquila_stLFR to reconstruct LFRs (Supplementary Figure 3). Ideally, each individual LFR is attached to the same unique barcode and sequenced by short-read sequencing, so short reads with the same barcode could be linked together to form the original LFR (linked-read). Aquila_stLFR reconstructs all LFRs relying on this concept. However, it is still necessary to tackle the barcode deconvolution problem: each group of reads that share the same barcode is drawn from an unobserved set of fragments, since the ideal design of one unique barcode per LFR will not be achieved for all LFRs during preparation of real sequencing libraries. Aquila_stLFR applies an empirical boundary threshold (50kb) to differentiate between two LFRs with the same barcode. For instance, if the distance between two successive reads with the same barcode is larger than 50kb, they will be assumed to be drawn from two different LFRs.

After reconstructing all LFRs, Aquila_stLFR applies a recursive clustering algorithm to partition LFRs into haplotype-specific blocks (Zhou *et al*., 2021). Aquila_stLFR further cuts large haplotype-specific blocks into small chunks (100kb by default). Finally, short reads that are drawn from LFRs are identified and reassembled locally by SPAdes (Bankevich *et al*., 2012) within each small chunk. To achieve large contiguity, Aquila_stLFR iteratively concatenates assembled minicontigs from small chunks into full-length contigs. To detect SVs from diploid assembly, Aquila_stLFR performs a pairwise comparison between haploid contigs and reference by integrating Minimap2 (Li, 2018) and paftools (https://github.com/lh3/minimap2/tree/master/misc). Additionally, the “Aquila_hybrid” mode applies an analogous concept to reconstruct long DNA fragments and generate the same data structure for long DNA fragments of both 10X and stLFR data (Supplementary Figure 1). Aquila_hybrid can combine both linked-read data for diploid assembly.

## 3 Results and Discussion

In this study, we used the stLFR sequencing library for NA24385 from the Genome in A Bottle (GiAB) website (ftp://ftp-trace.ncbi.nlm.nih.gov/giab/ftp/data/AshkenazimTrio/HG002_NA24385_son/stLFR/), since we can evaluate SV (≥ 50bp) detection by comparing Aquila_stLFR’s calls against the GiaB NA24385 SV benchmark callsets. This stLFR library has approximately 100x Illumina sequencing coverage, and the average inferred (reconstructed) DNA fragment (LFR) lengths are around 26kb. To demonstrate the performance of SV calling for Aquila_stLFR, we have compared it against two other linked-reads SV callers: GROC-SVs and NAIBR (Table 1). After generating SV calls for NA24385 from all three methods, we used Truvari (parameters −p 0.1, −P 0.1, −r 200, −passonly) to compare SV calls with the benchmark callset from three different SV size range thresholds. In table 1, we show that Aquila_stLFR achieved a higher sensitivity than the other two methods of 80.5% for deletions SV ∈ [50bp, 1kb] and 63.5% for deletions SV ∈ [1kb, 10kb]. NAIBR achieved a higher sensitivity of 82.8% for deletions (SV > 10kb). Since few insertions were found by GROC-SVs and NAIBR, we only performed insertion evaluation for Aquila_stLFR. Aquila_stLFR achieved a high specificity of 83.8% with a sensitivity of 23.4% for insertions SV ∈ [50bp, 1kb]. In conclusion, Aquila_stLFR is the first approach that effectively leverages the strengths of linked-reads/Illumina sequencing to enable medium to large SV discovery.

**Table 1.**
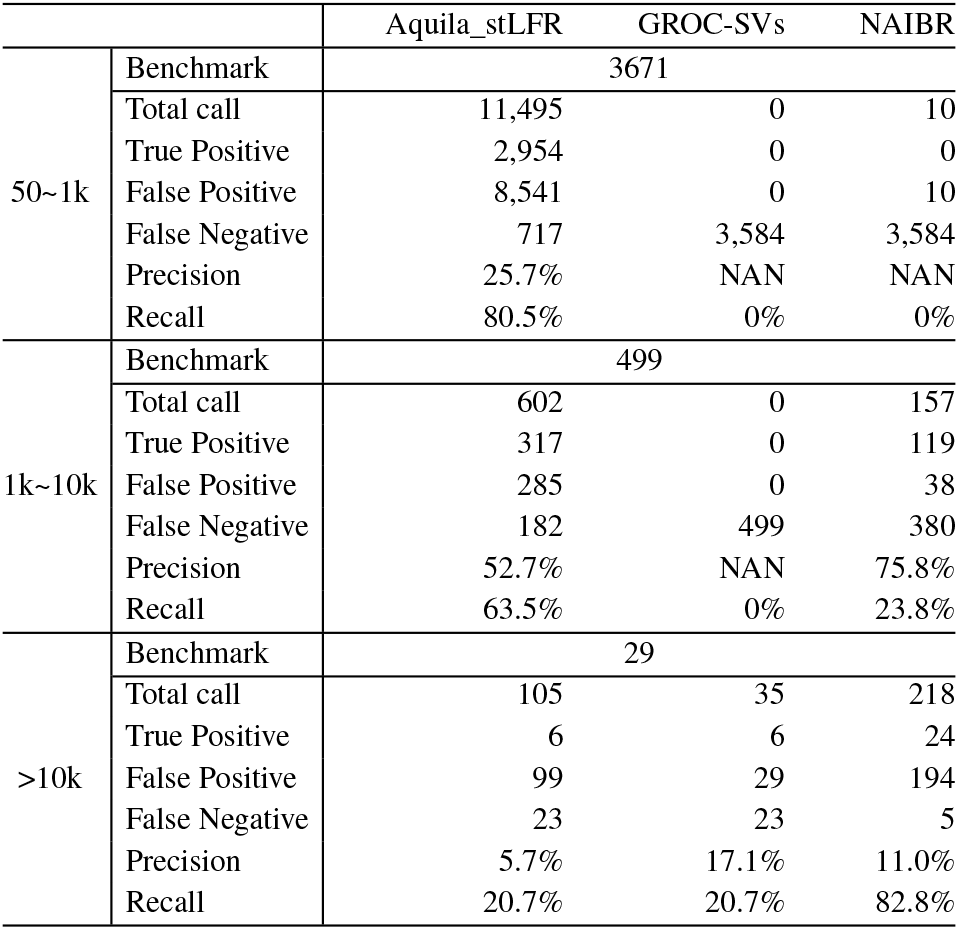
Genome-wide deletion evaluation against GIAB NA24385 benchmark.

Our study also provides a guideline for future stLFR library preparation to achieve significant improvement in assembly and SV detection. Recent studies with different customized 10X linked-reads libraries showed that the optimal physical coverage *C_F_* was between 332X and 823X. They also suggested that the optimal length-weighted fragment length (*μ_FL_*) was around 50 - 150kb (Zhang *et al*., 2019, 2020). We found that the stLFR library had a configuration of these three parameters (*C_F_* = 238X, *C_R_* = 0.35, *μ_FL_* = 26kb). The relatively low *μ_FL_* was one key cause of the lower contiguity of stLFR assemblies compared with that of 10x linked-reads assemblies, which then influenced SV calling. Furthermore, the stLFR library used 100bp paired short reads which created a limitation for local assembly compared to the 150bp paired short reads used by 10X linked-reads. Future studies may take advantage of the tools and analysis we present here.

## Supporting information

Aquila_stLFR_SupplementaryInformation

## Acknowledgements

This research was supported by Vanderbilt University Development Funds (FF_300033), the Joint Initiative for Metrology in Biology (JIMB; National Institute of Standards and Technology) and Research Grant Council Early Career Scheme (HKBU 22201419).

## Notes

### Competing Interest Statement

The authors have declared no competing interest.

### Summary of Updates

We have revised the paper to a short paper version and reordered the authorship. We will submit it to a journal later.

