## Supplementary material for "Aquila_stLFR: diploid genome assembly based structural variant calling package for stLFR linked-read": Aquila_stLFR_SupplementaryInformation

### Supplemental Material

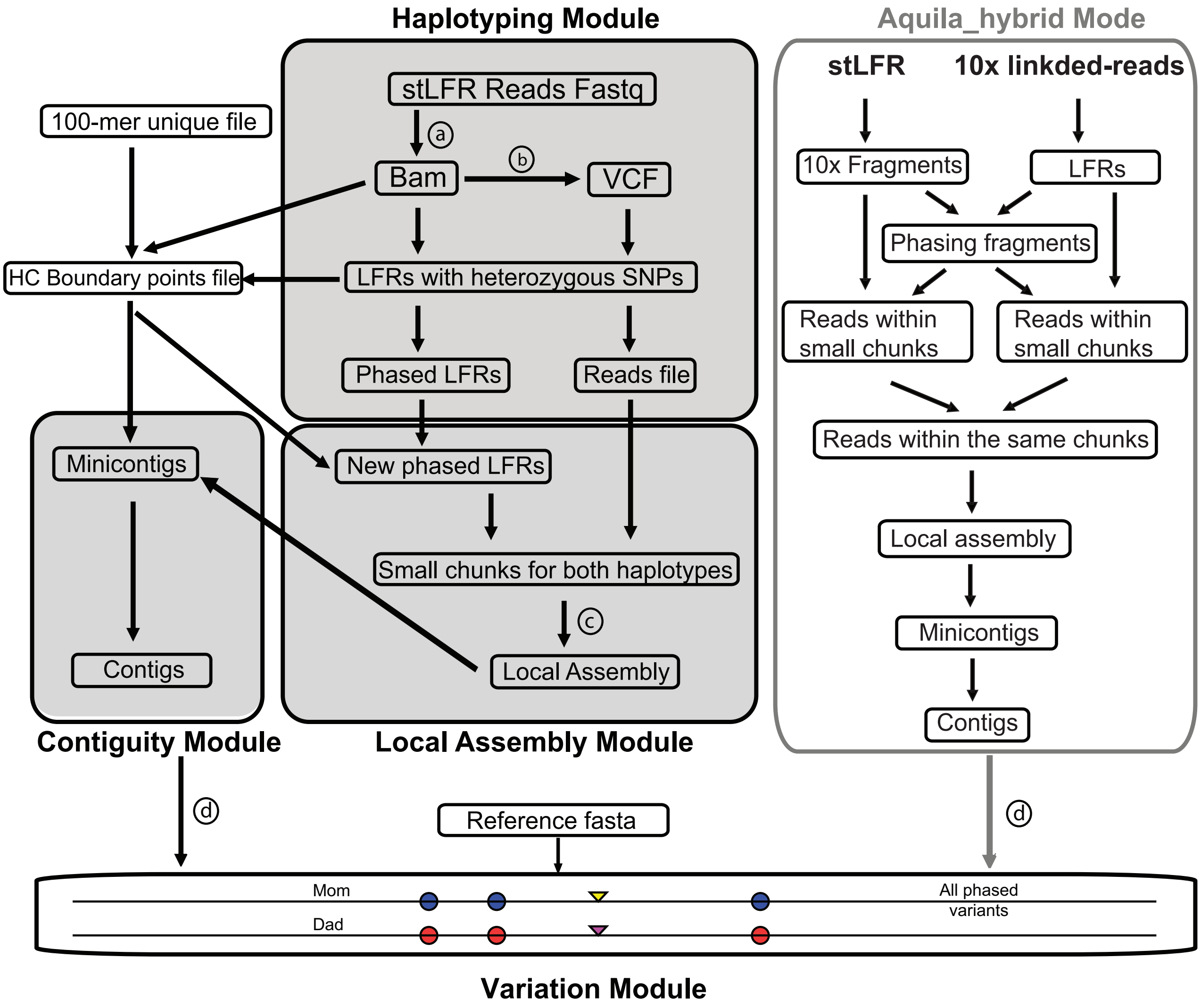


Supplemental Figure 1: Pipeline of Aquila_stLFR, a reference-assisted diploid-resolved genome assembly for stLFR. Input files: FASTQ file, BAM file and VCF file. a: Bwa-mem; b: FreeBayes; c: SPAdes; d: minimap2 and paftools.

```
@CL100066606L2C016R059_503217   BX:Z:540_839_548
AAAACACACTTTTATTTTATTTTATTTTATTTATTTAATTTAATTTAATTTAATTTTATTTTATTTTATTTATTTTATTTTATTTTATTTTATTTTTTTG
+
A@;,<EEEE<6B=EEDAFEEEC@FECE8DE@BB=ECCCAFBA-E?ABD>BEF3EE=@332E@EB>E.EABE:D=EDAA6ECFEBE=AEEFFB@EBE?>E9
@CL100066606L2C016R059_503217   BX:Z:540_839_548
CCTGTAGTCCCAGCTACTCAGGAGGCTGAGGCAGGAGAATGGCGTGAACCCAGGAGACAGAGCTTGCAGTGAGCTGAGATAGCGTCACTGCACTCCAGCC
+
EEFFE<FEDEFFFEFEEEF?EEAF8FFD:?EFEFEAD2D6<EFEFF@FFEEFED?EAEDF3DFFFFECFFE,EFFFAFEFDCFEFFDCFFCDFFFD=<<A
@CL100066606L2C016R091_430379   BX:Z:540_839_548
CTCACCCCTTCTACCATGTGAGGACGCAGCAAGAAGTCACCTTCTATAAATCAGGAAGAGATCGCTAACCATCCCCTGTATCTGCTAGTGCCTGGATCTT
+
F5F>EFFFD*FA<FED;F>DCFF==FFDEFFFFA=F8FFFF:EFAE4E-0<7<?FA<FBF'@AFF3B/=FA@FFF<=F(ECF8FF5FFAFE8@FF?DF-=
@CL100066606L2C016R091_430379   BX:Z:540_839_548
CATATAACCTCACCCACATAGAGAAATAAAATAACTAGGATCCAGATGTTCCTGCATAAACCCTCTTGATTCTTACCCATTCACATTCTCCAGAAATAAC
+
CFDEAB6FDFEEFDE<F>4AE4=+4@FBD;:BE>EE;EE29AF:F1DF5/EE8'E;6<5%CEECBB.F0>3EBBFB;F>A3E?F;C.8<BF2&&@9;E,E
@CL100066606L2C017R026_292022   BX:Z:540_839_548
CCTTCACTTTCAGGTTTTAATGTATTCAACTCATTTGGATAAATACCAAGGAGCACAGCCTCTTCTTTTTTATTATTGAGCAATCTCAAATGCATCTCTG
+
FEEEFF.EF<FFFDEEF@E@@EFFCBFD?EEECEDAFBFCD@FEFF?F:E>?FEFD>3EFCCE455FDCE?F9DEE>D8:EF8DEFEEFEEFF=6FEFDE
@CL100066606L2C017R026_292022   BX:Z:540_839_548
CCCTCCTTTGGGAGATGCTGCTCTTGAAGTAGAAGAAAAGTGCTAACTGGAAATTCTTTGAAACCTCACTTTCTCATTGTCAGCTGGTCAAAAGAAAACA
+
FFFFF>EFFFFDFFFFFFEFFFFFEFEE:EFFFFFCFFDFFFFFF@FFFEC56EFFFDDF>9=FFFFFFFE@<FF'E>FBFCFFDGFE,F/@@E=D4EFF
```

Supplemental Figure 2: A screen shot of stLFR fastq reads, it shows three pairs of short reads. Before performing reads alignment by bwa-mem, add barcode “BX:Z:barcode” at the header of each read.

```
CL200047468L1C006R038_39389     147     chr1    9997    0       100M    =       10013   -84     CAGATAACCCTAACCCTAACCCTAACCCTAACCCTAACCCT
AACCCTAACCCTAACCCTAACCCTAACCCTAACCCTAACCCTAACCCTAACCCTAACCC     9CFEFGFAFG7FGFEFEGFDFFG6E@FFG?FFFFGDGFFFFFFFFFGEFFFFFGFEFFGG@FF@GGFFFFGGF
FF?FGDFFGGFFFFFEGFG;FFFA<FF     NM:i:1  MD:Z:1C98       AS:i:98 XS:i:96 BX:Z:586_902_1136       RG:Z:12878:LibraryNotSpecified:1:unknown_
fc:0-43F6A2D2
CL200047469L1C011R060_87815     147     chr1    9997    0       100M    =       10013   -84     CAGATAACCCTAACCCTAACCCTAACCCTAACCCTAACCCT
AACCCTAACCCTAACCCTAACCCTAACCCTAACCCTAACCCTAACCCTAACCCTAACCC     FFDFFFCGFGEFGFFFFG1FFGFCFFFFGFGFFGE<FFFFFFGFFFGFGFFFEGG>FFFGFFFFFFFFFFFFF
FFFEFF>FFGFFFFFEFFFFFFFFFFF     NM:i:1  MD:Z:1C98       AS:i:98 XS:i:96 BX:Z:353_1476_263       RG:Z:12878:LibraryNotSpecified:1:unknown_
fc:0-64378858
CL100066606L2C006R022_551816    147     chr1    9997    0       100M    =       10007   -90     CAGATAACCCTAACCCTAACCCTAACCCTAACCCTAACCCT
AACCCTAACCCTAACCCTAACCCTAACCCTAACCCTAACCCTAACCCTAACCCTAACCC     FFFECDDFFFFFFDEFE>FECFFE6FEDFFGFEFF>FDBFFCFFFDFFFFDFFFFFFFFE6FFFFFFFCEFFF
FDEFFFFFFFFFFFDGFFEFFFFFFEC     NM:i:1  MD:Z:1C98       AS:i:98 XS:i:96 BX:Z:0_0_0      RG:Z:12878:LibraryNotSpecified:1:unknown_fc:0-43F
6A2D2-5F69628
CL200047469L1C009R005_5964      163     chr1    9998    0       100M    =       10005   108     CCATAACCCTAACCCTAACCCTAACCCTAACCCTAACCCTA
ACCCTAACCCTAACCCTAACCCTAACCCTAACCCTAACCCTAGCCCTAACCCTAACCCT     5E&F=?.)<@E1BD2DCF7*76FE<2<.E)6<-54<)(4?<-7D75F5BEED>>8=BEEF83;8?E4A0C-AA
*8=E1A,ECB%:1AFFC7F@BC03EDB     NM:i:2  MD:Z:1G81A16    AS:i:93 XS:i:92 BX:Z:46_1232_9  RG:Z:12878:LibraryNotSpecified:1:unknown_fc:0-515
43204
CL100066606L2C017R018_461042    99      chr1    9998    0       100M    =       10044   146     CCATAACCCTAACCCTAACCCTAACCCTAACCCTAACCCTA
ACCCTAACCCTAACCCTAACCCTAACCCTAACCCTAACCCTAACCCTAACCCTAACCCT     >:)GG@EFFGFD0F?EGFFGGEFFGFGBFFCGFBGCFFFFFEEC<GFB=FFFFFF7DFGGFDBFGDFDGEFFD
GGBE@FFEDF5:FGFG@BFGB7EFGFF     NM:i:1  MD:Z:1G98       AS:i:98 XS:i:97 BX:Z:584_924_493        RG:Z:12878:LibraryNotSpecified:1:unknown_
fc:0-64717E9D
```

Supplemental Figure 3: A screen shot of five reads from BAM file. Each read contains the barcode field “BX:Z:barcode”.
